# ChIP-AP – An Integrated ChIP-Seq Analysis Pipeline

**DOI:** 10.1101/2021.04.18.440382

**Authors:** Jeremiah Suryatenggara, Kol Jia Yong, Danielle E. Tenen, Daniel G. Tenen, Mahmoud A. Bassal

## Abstract

ChIP-Seq is a technique used to analyse protein-DNA interactions. The protein-DNA complex is pulled down using a protein antibody, after which sequencing and analysis of the bound DNA fragments is performed. A key bioinformatics analysis step is “peak” calling - identifying regions of enrichment. Benchmarking studies have consistently shown that no optimal peak caller exists. Peak callers have distinct selectivity and specificity characteristics which are often not additive and seldom completely overlap in many scenarios. In the absence of a universal peak caller, we rationalized one ought to utilize multiple peak-callers to 1) gauge peak confidence as determined through detection by multiple algorithms, and 2) more thoroughly survey the protein-bound landscape by capturing peaks not detected by individual peak callers owing to algorithmic limitations and biases. We therefore developed an integrated ChIP-Seq Analysis Pipeline (ChIP-AP) which performs all analysis steps from raw fastq files to final result, and utilizes four commonly used peak callers to more thoroughly and comprehensively analyse datasets. Results are integrated and presented in a single file enabling users to apply selectivity and sensitivity thresholds to select the consensus peak set, the union peak set, or any sub-set in-between to more confidently and comprehensively explore the protein-bound landscape. (https://github.com/JSuryatenggara/ChIP-AP).

## Introduction

ChIP-Seq is an extensively used experimental technique that aims to identify DNA binding location sequences and motifs of DNA-interacting proteins such as transcription factors^1,2^, histone-modifier proteins^3^, or novel DNA-binding proteins. To perform a ChIP-Seq experiment, cells are fixed, the chromatin-protein complex sonicated, and the DNA fragments interacting with the protein of interest pulled down by the targeted protein antibody. Following experimental pull-down of the protein-bound DNA and sequencing, the raw sequencing data (raw fastq files) undergoes processing and analysis, after which, biological relevance can be inferred^4^. Such analyses, in conjunction with follow up mechanistic studies, shed light on the DNA-associated proteins biological function and roles^4^.

Since the development of ChIP-Seq^2,5,6^, computational analysis of ChIP-Seq experiments has always been a multi-step process requiring multiple command line programs which, to use most effectively, requires knowledge and experience in computing and programming^4^. Wet-lab biologists without command line or coding experience have typically relied on bioinformaticians, with their computing expertise, to analyse the sequenced data despite them potentially having a reduced understanding of the underlying biology. Irrespective of whom performs the analysis however, in the computational space, many analogous programs have been developed for each stage of the analysis, complicating how one should approach an analysis and the decision of which programs to use in conjunction with one-another. It is therefore easy to understand why two different analysis methodologies, even if they appear superficially identical or similar, will almost certainly report different results, leading to conflicting conclusions from the same biological experiment^7,8^. To complicate matters further, published methods outlining the workflow used to analyse a dataset will consistently lack essential details, with some authors omitting key program modification parameters/flags, or neglecting to include key analysis steps entirely, relegating published analyses to being almost entirely irreproducible for other researchers.

Of all the complications plaguing ChIP-Seq analyses though, perhaps the most well-known source of inconsistency between analyses is the choice of peak calling algorithm^2,7,8^. This observation was convincingly demonstrated by Steinhauser et. al.^8^ in their study comparing 20 peak callers wherein they reported poor agreement between the called peak sets across the profiled callers. This, and other studies, therefore show that peak callers have distinct selectivity and specificity characteristics which are often not additive and seldom completely overlap in many scenarios^8–11^. Consequently, such differing operating characteristics results in a lack of consistency across the reported regions of enrichment (and associated genes) for each peak caller. This has follow-on effects in that downstream functional analysis for the protein of interest would therefore give differing, potentially conflicting results. Additionally, it has been shown extensively that the performance of a peak caller is subject to the read characteristics and read distributions of the dataset in question^8–12^. An individual peak caller can outperform other callers in certain datasets but will perform poorly in alternate datasets. Therefore, relying on a one-caller-fits-all approach when analysing datasets with different DNA-binding proteins, immunoprecipitation and library preparation protocols is objectively not the soundest approach to yield reliable, consistent and comprehensive results.

We therefore rationalized that in order to improve the reliability and consistency of a ChIP-Seq analysis, one ought to focus on and improve the consistency, confidence and comprehensiveness of the peak detection step without requiring additional wet-lab observations. To address this, we designed our ChIP-Seq Analysis Pipeline (ChIP-AP) which integrates all processes of a ChIP-Seq analysis (from raw fastq to final result) into a single, easy to use package, that utilizes four commonly used peak callers^13–17^ (for either transcription factors or histone modifier proteins), and integrates their results into a single output file, from which, users are able to infer peak confidence. If a peak is called by multiple callers, one can infer that the reported peak has a higher confidence and is less likely to be a false-positive or an artefact of algorithmic bias or limitation. Alternatively, by integrating the results of all the callers (the union of all peaks), one can more comprehensively survey the binding landscape of the binding protein by capturing peaks that would otherwise be uncalled or “lost” if relying on a lone peak caller. In other words, the union peak set enables a more comprehensive survey of the binding landscape by accepting all peaks irrespective of confidence. By utilizing multiple peak callers and integrating their results, users are able to determine the selectivity and specificity requirements that best describe their dataset while allowing them to circumvent inherent sample characteristics that result in poor peak calling performance of any single peak caller, thus enabling the capture of either the most number or peaks (the union of all peaks), the most confident peaks (the consensus results), or any sub-set of the gradient in-between. ChIP-AP can therefore become an effective tool for users by providing both substantial improvements to peak capturing and analysis reliability. ChIP-AP is available on GitHub (https://github.com/JSuryatenggara/ChIP-AP) with extensively detailed wiki pages describing installation, use and results interpretation (https://github.com/JSuryatenggara/ChIP-AP/wiki).

## Results

### Consensus peaks increase motif and ontology accuracy

A reproducible result instils greater confidence in its validity. Likewise, ChIP-Seq peaks that can be detected by multiple peak callers, each utilizing different peak detection algorithms, garner greater confidence than peaks called by an individual peak caller. ChIP-AP, with its utilization of four different peak callers, reports along-side the coordinates of a peak how many callers detected the said peak. Using this, users are able to filter on the consensus peaks, which are the peaks detected by all four peak callers and, consequently, carry the greatest confidence. We hypothesized that utilizing the consensus peak set would increase the percentage of peaks containing a valid binding motif (peak-motif percentage) without drastically affecting the motif position bias (the distance of the motif to the weighted peak center). The consensus peak set would also increase the likelihood of identifying correct binding motifs while masking co-factor binding artifacts. Finally, the consensus peak set should also improve gene ontology (GO) results by ensuring only the strongest binding candidates are included in analysis.

To investigate whether the peak-motif percentage for the consensus peak set is significantly better than using a single peak caller (MACS2), we processed 10 transcription factor (TF) datasets from differing TF families, across 3 cell lines sourced from ENCODE^18^, and determined their peak-motif percentage (see **Methods** for sample details). For each TF, we downloaded the binding motif from MethMotif^19^ (which sourced its motifs from JASPAR 2018^20^), and determined how many peaks contain the binding motif while allowing up to a single sequence mismatch. As expected, a significant difference in the average peak-motif percentage was observed across all 10 TF’s (two-tailed t-test p=0.0012, **Figure 1a**). The degree by which the consensus peak set improved the peak-motif percentage was variable, with an up to 90% improvement for RUNX1, but none the less, still showed improvement for all 10 TFs over the MACS2 peak set alone (**Supplemental Table 1**).

**Figure 1.**
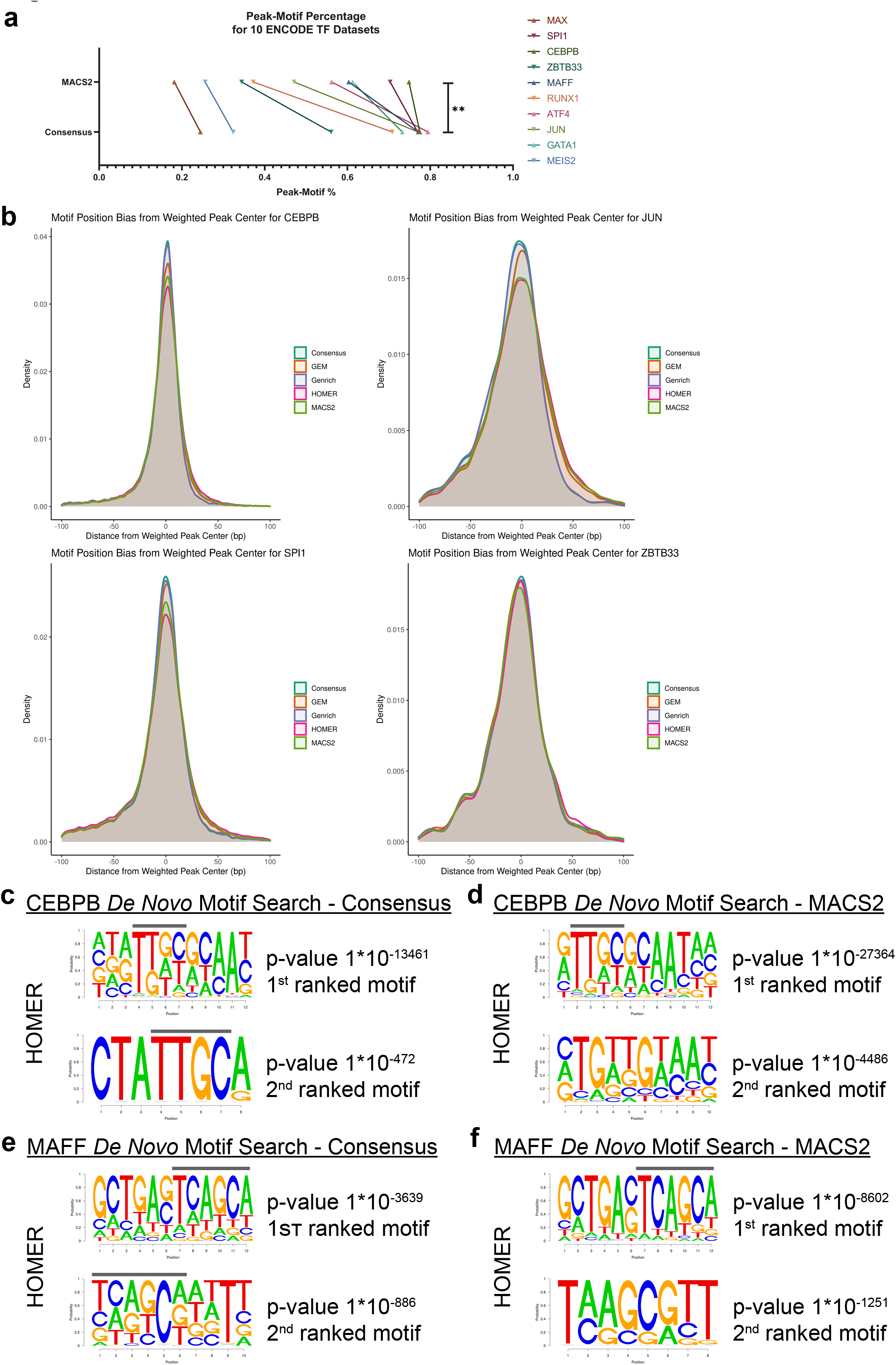
Consensus peak set improves detected motif accuracy. a) Peak-Motif percentage (number of peaks with binding motif) for all 10 TF’s profiled as identified for the MACS2 and the consensus peak sets. b) The motif position-bias for CEBPB, JUN, SPI1 and ZBTB33 for the consensus peak set and the individual peak callers. The position-bias is a measure of how far the identified motif sits away from the weighted peak center. c) The CEBPB *de novo* motif discovery results as reported by HOMER for the consensus peak set. The line above the peaks delineates position of the binding motif. d) The CEBPB *de novo* motif discovery results as reported by HOMER for the MACS2 peak set. The line above the peaks delineates position of the binding motif. e) The MAFF *de novo* motif discovery results as reported by HOMER for the consensus peak set. The line above the peaks delineates position of the binding motif. f) The MAFF *de novo* motif discovery results as reported by HOMER for the MACS2 peak set. The line above the peaks delineates the position of the binding motif.

We next investigated whether the consensus peak set significantly altered the motif position bias with respect to the weighted peak center as compared to individual callers. Across all 10 TFs, the benefit of the consensus peak set was variable, with it out-performing all individual peak callers in some datasets (**Figure 1b, Supplemental Table 2**), while in others, providing comparable results (**Supplemental Figure 1a, Supplemental Table 2**). Consistently though, the consensus peak set did show significant improvement over at least half the peak callers tested suggesting that utilizing the consensus peak set will either give an improved motif position bias profile or report comparable results to having used an individual peak caller.

Next, we questioned whether the consensus peak set can provide improved *de novo* motif sequence detection while masking co-factor binding artefacts. Previously, Lin et al.^21^ described that *de novo* global motif analyses can potentially be contaminated by co-factor motif sequence artefacts. In their publication, they highlighted potential co-factor motif artefacts for CEBPB and MAFF. Using ChIP-AP, we performed *de novo* motif analysis using HOMER^15^ and the MEME-Suite^22–24^ for the MACS2 called and the consensus peak set for these two TFs. For CEBPB, the first candidate motif hit reported by HOMER for both the consensus and the MACS2 peak sets was near identical, which, according to Lin et al.^21^, contains co-factor binding motif artefacts (**Figure 1c, d** upper panels).

However, the second motif hit for the consensus peak set (**Figure 1c** lower panel) shows the clean binding motif sequence TTGC, which is the CEBPB motif that contains neither co-factor motif contamination nor a heterodimer sequence^21^. The MACS2 peak set’s second motif result however (**Figure 1d**, lower panel), failed to report the same motif result. When analysed using the MEME-Suite, the consensus peak set showed two motifs with co-factor artefacts (**Supplemental Figure 1b**, 1^st^ and 3^rd^ ranked), two motifs with the heterodimer sequence (**Supplemental Figure 1b**, 2^nd^ and 4^th^ ranked) and the fifth result showed the TTGC binding motif without co-factor motif contamination nor a heterodimer sequence. Conversely, for the MACS2 dataset as analysed by the MEME-Suite, three motif results contain co-factor sequence artefacts (**Supplemental Figure 1c**, 1^st^, 3^rd^ and 4^th^ ranked), one result with heterodimer sequence (**Supplemental Figure 1c**, 2^nd^ ranked) and the fifth result was also the TTGC binding motif without co-factor motif contamination nor a heterodimer sequence. Therefore, for CEBPB, although both *de novo* motif algorithms reported similar findings, the consensus peak set showed a cleaner and more direct signal for the CEBPB binding motif from the second HOMER result, a finding not immediately evident in the MACS2 set without careful inspection of the data.

For the TF MAFF, the HOMER *de novo* motif result for the consensus peak set remained more consistent (**Figure 1e**) with both the top two motif results showing the binding sequence TCAGCA. The MACS2 peak set however, wasn’t as consistent showing different sequences between the first and second motif candidates (**Figure 1f**). Similarly for the MEME-Suite results (**Supplemental Figure 1d, e**), MEME consistently calls the TGCTGA for both peak sets but the 6^th^ reported motif candidate for the consensus peak set shows a heterodimer binding profile (characterised by the “sequence - spacer - reverse complement sequence” profile), a result not recapitulated in the MACS2 peak set entirely, thereby supporting the notion that the consensus peak set can provide more direct *de novo* enriched motif results over using a single peak caller alone.

Our final investigation was to test whether the consensus peak set can provide improved, more direct gene ontology (GO) results. In running a GO analysis for all 10 TF’s profiled and comparing the consensus peak set results to the MACS2 peak sets, we observed that for certain datasets, such as RUNX1, ATF4, JUN, ZBTB33 and GATA1, the consensus peak set GO results returned more relevant and directly related terms than the MACS2 peak set (**Table 1, Supplemental Tables 3, 4**). For the RUNX1 results, whereas the top 20 MACS2 GO results contained generic GO terms, the consensus peak GO listing clearly outlined RUNX1 functions regarding hematopoietic differentiation, regulation of metabolic and signalling pathways and autophagy regulation, all of which are known published functions of RUNX1^25–30^ (**Table 1**). The GO terms returned when searching the consensus peak candidate list can therefore, for certain datasets, provide significantly clearer and more direct GO results by providing information on only the gene terms corresponding to the most confident peaks called by all peak callers.

Therefore, utilizing the consensus peak set can provide added benefits to identify novel binding motifs or to identify more direct biological processes modulated by a protein of interest, especially if it is not well characterized as evidenced by the results presented. In all metrics investigated, the consensus peak set’s performance was either significantly improved, or, in worst performing cases, provided results comparable to having used only a single peak caller.

### Capturing Lost Peaks with the Union Peak Set

A number of variables can affect a ChIP-Seq experiments efficiency resulting in poor enrichment and potentially giving rise to a high signal:noise ratio dataset. Every peak caller has differing operating characteristics and thus, has differing abilities to handle these difficult to process datasets^8,12^. A ChIP-Seq analysis utilizing only a single peak caller would be solely dependent on the chosen peak callers’ ability to handle the signal:noise ratio and enrichment characteristics of that dataset. If the peak caller struggles to differentiate signal from noise effectively, few peaks will be called and a dataset will give an inconclusive result owing to its ineffectiveness to deal with the dataset. However, some peak callers are more capable at handling difficult datasets, and so an experiment may show poor enrichment, but it simply needs to be analysed with the right peak caller for its specific characteristics, the choice of which may not be evident or obvious in advance.

One protein that is relatively difficult to perform ChIP-Seq on, is the oncogene sal-like protein 4 (SALL4). SALL4 has been shown to play essential roles in maintaining pluripotency and self-renewal characteristics of embryonic stem cells (ESC)^31^. It is typically down-regulated after birth but has been found to be aberrantly regulated in many tumors^31,32^. Studies have also shown SALL4 to have multiple protein interacting partners and DNA-binding and regulation functions^31,33^. An attempt to capture the DNA-binding partners of SALL4 was undertaken with the sequenced result showing poor enrichment on the fingerprint plot with little separation between the SALL4 ChIP-Seq replicate and control curves (**Figure 2a**), indicating it will likely be difficult to call peaks for this dataset. When processed with ChIP-AP, we observed that peak callers GEM and MACS2 struggle to call peaks (**Figure 2b**) with each returning a total of 1,362 and 1,937 peaks respectively. HOMER, is able to call approximately double the number of peaks at 3,760. However, Genrich, which determines peaks using an area under the curve (AUC) calculation rather than generating a Poisson distribution model (as seen in MACS2, GEM and HOMER), is more successful in dealing with such a dataset and calls a total of 12,452 peaks. We therefore sought to investigate the efficacy of utilizing the union peak set for this poorly enriched SALL4 ChIP-Seq, which enables us to sacrifice specificity for a gain in sensitivity across the dataset, ie, we accept all peaks including those called by only a single peak caller which carry less confidence but provide higher sensitivity (**Supplemental Table 5**).

**Figure 2.**
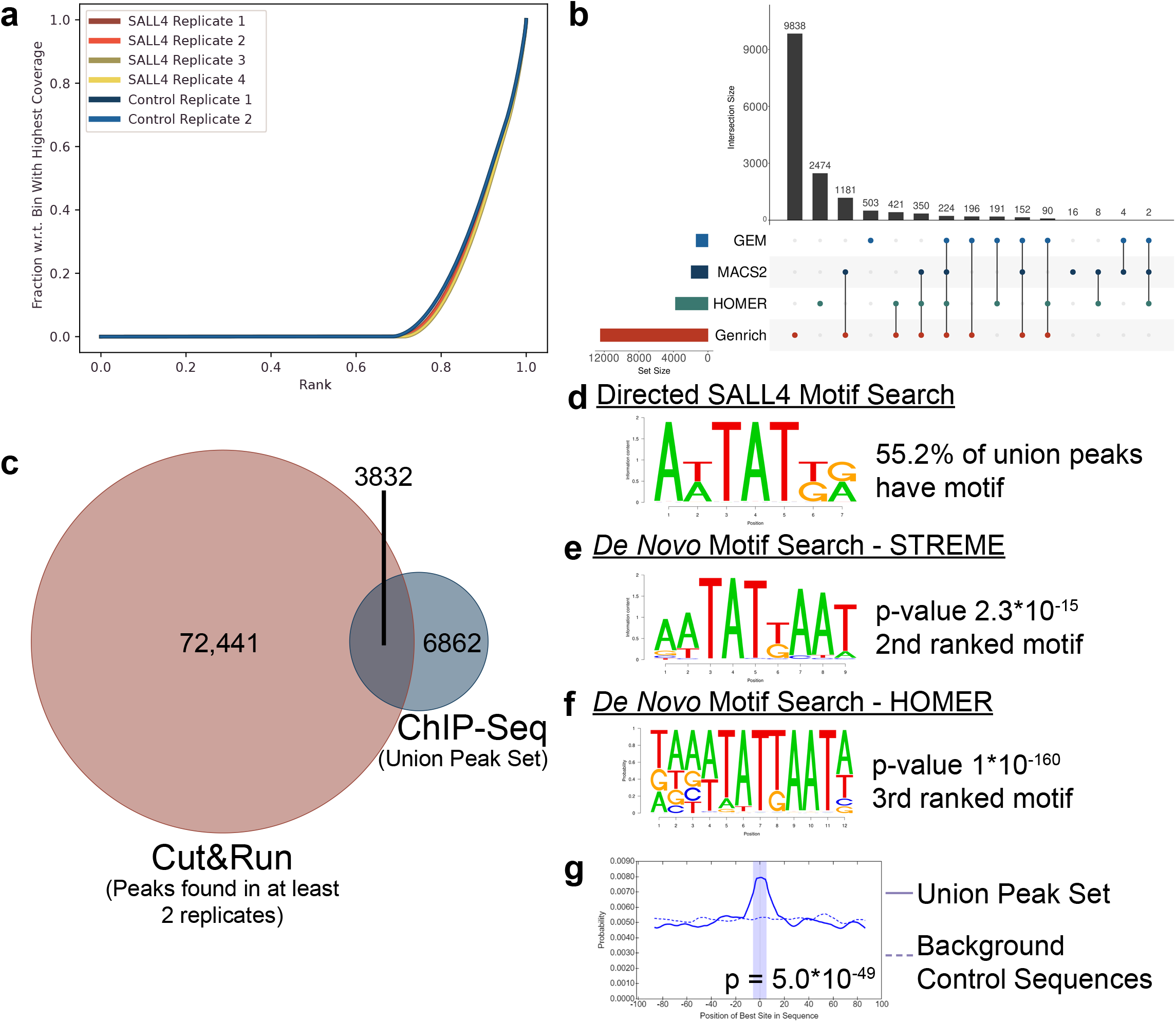
Union peak set comprehensiveness and accuracy. a) Fingerprint plot for aligned sequence files for samples. Negligible separation between the SALL4 and control curves indicates poor enrichment in the SALL4 samples. b) Upset plot describing the distribution of peaks observed by each peak caller. The left histogram represents the total number of called peaks per caller. The top histograms represent the size of the sub-sets in question. The connected circles represent highlighted overlap. c) Venn diagram showing the overlapping number of peaks between the SALL4 union ChIP-Seq dataset and the Cut&Run dataset. d) The motif sequence used for the directed motif search in the SALL4 ChIP-Seq union set, which was found in 55.2% of the union set. e) The STREME *de novo* motif search for the SALL4 union peak set identified the AT-rich binding motif as the 2^nd^ result. f) The HOMER de novo motif search for the SALL4 union peak set identified the AT-rich binding motif as the 3^rd^ result. g) The STREME identified motif (shown in d) was found centrally enriched in the union peak set as compared to background sequences.

To test its validity, we compared the SALL4 ChIP-Seq union peak set with a SALL4 Cut&Run dataset recently published^33^. Cut&Run is a technique which utilizes antibody-targeting and micrococcal nuclease digestion to map global DNA binding sites^34^. It is an analogous but independent technique to ChIP-Seq thus providing an independent dataset for comparison and validation. To ensure the peaks called in our ChIP-Seq were likely binding targets of SALL4, we first directly compared the union peak set to the SALL4 Cut&Run dataset. Reassuringly, the union peak set showed a 36% overlap with peaks identified in at least 2 of the Cut&Run replicates (3 biological replicates total) (**Figure 2c**). Using individual peak callers, overlap percentages ranging from 25-56% were observed with fewer peaks called (**Supplemental Figure 2a**). Furthermore, each caller reports a different sub-set of targets with little overlap between them (**Figure 2b**). However, by considering the union peak set, we can gain a more complete overview of the binding landscape without significantly affecting average sensitivity, by allowing us to circumvent the poor performance of individual peak callers for the dataset in question and call “missed” peaks.

Next, we wanted to confirm that the called union peaks show our recently identified human SALL4 DNA binding motif^33^. To investigate this, we performed a directed motif search wherein we searched every peak in the union peak set for the human SALL4 DNA binding motif. This showed that 55% of the union peak set contained at least one instance of our identified SALL4 motif (**Figure 2d**), a result comparable to using an individual peak caller alone (**Supplemental Table 1, 6**). To ensure we have not biased the motif search, we performed a *de novo* motif search on the union peak set using both the MEME-Suite^22–24^ (which utilize the algorithms STREME, CentriMo and MEME-ChIP) and HOMER which were both able to call the human SALL4 DNA binding motif as the second and third top candidate motif hits respectively (**Figure 2e, f**). According to CentriMo, the STREME identified motif is centrally enriched in individual peaks in the union peak set (**Figure 2g**), an expected observation for true binding motif sequences. Furthermore, MEME-ChIP itself, called the same motif as the third candidate peak with the second candidate motif result also being an AT rich motif with near identical sequence (**Supplemental Figure 2b**). We therefore concluded that despite the additional number of peaks called by taking the union peak set, the SALL4 DNA-binding motif signature is still present across all called peaks and is identifiable using multiple algorithms as a top three candidate motif.

As further validation to ensure the union peak set identified valid targets of SALL4, a GO analysis was performed and compared with the results of previous findings^33^. We previously reported that SALL4 knock-down resulted in a significant increase in the number of up-regulated genes in the “*transcriptional regulator activity*” (GO:0140110) pathway, results which were validated by comparing bulk RNA-Seq and Cut&Run results^33^, and thus confirming pathway members as *bona fide* SALL4 targets. Consistently, the GO analysis on the union peak set identified the same pathway as a top 20 enriched pathway (**Supplemental Table 7**), with more significantly enriched terms pointing to SALL4 being a DNA-binding protein; a well-established function of SALL4^31,32^ (**Supplemental Table 7**). Therefore, despite utilizing the union peak set which sacrificed a degree of specificity, the peak set was still valid in detecting accurate biological functions of SALL4.

The final validation to ensure the union dataset identified valid targets of SALL4 was to overlap the union peaks gene list with the SALL4 knock-down bulk RNA-Seq previously published^33^, and compare the overlap targets of the union peak set with the overlapping targets of the Cut&Run dataset. We previously reported that 2,695 genes were found significantly differentially expressed on SALL4 knock-down, 430 of which has a corresponding annotated SALL4 Cut&Run peak. Using the union peak set, we observed an overlap of 451 genes (**Supplemental Table 8**) with the SALL4 knock-down dataset, of which, 198 gene targets were found in common between the Cut&Run and ChIP-Seq gene sets (**Supplemental Table 8**). This finding combined with the observed overlap between the union peak set and the Cut&Run replicates suggests that there are SALL4 binding targets that were detected by the ChIP-Seq that were not detected by the Cut&Run and vice versa. However, both the Cut&Run and the union peak list derived from the SALL4 ChIP-Seq appear to still be calling valid SALL4 target genes with significant overlaps between the two datasets observed.

Taken together, the results obtained show that despite this SALL4 ChIP-Seq showing poor enrichment with few peaks called by two of the four peak callers, there was still valid data within the dataset that can be extracted, used, and validated by independent approaches^33^. By considering the union peak set generated by ChIP-AP, one can opt to marginally sacrifice specificity for a significant gain in sensitivity across the dataset and confirm the presence of peaks identified or validated using different methodologies should the characteristics of the dataset prove to be less than favourable. Whereas previous analyses using a single peak caller would produce sub-optimal results, by relying on multiple peak callers, as ChIP-AP does, sub-optimal datasets can be salvaged and still report valid findings.

### ChIP-AP Functionality and Characteristics

#### ChIP-AP Modularity for Advanced Users

Many programming languages are based on the programming paradigm of Object-Oriented Programming (OOP), wherein individual components of the program resemble “reusable objects” with defined input and output parameters (**Figure 3a**). This compartmentalization allows the programmer to assemble these “objects” in any manner to accomplish the task at hand provided the requisite parameters are met for individual objects. In the same spirit as OOP, ChIP-AP has been designed to be “object-oriented” in nature (**Figure 3b**).

**Figure 3.**
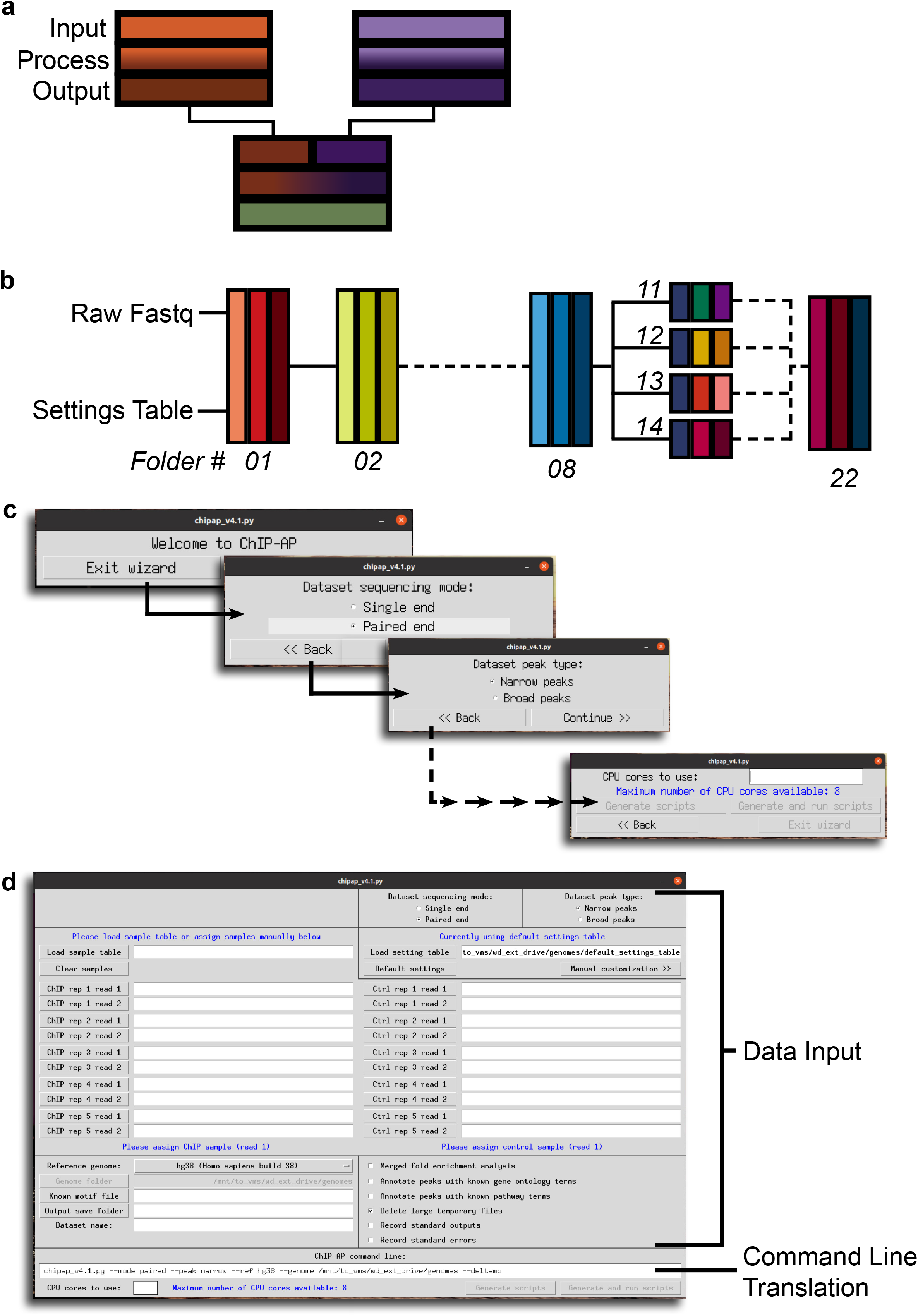
Object Oriented Nature of ChIP-AP. a) In OOP, an abstract “object” is defined as a segment of code that accepts defined inputs, processes the data, and outputs the data in a defined manner. Objects can then be combined in any manner to produce desired output. b) ChIP-AP was designed to be “object-oriented” in nature with each stage of analysis in a folder (01, 02…) having defined input/output characteristics. c) The ChIP-AP wizard interface guides users through a series of windows, each asking for a single piece of input, till all required information is gathered for a ChIP-AP run. d) The dashboard interface can be separated into 2 regions, the data input and command line translations segments. In the data input section, all the required data for a ChIP-AP run is input from a single interface. The command line translation window at the bottom dynamically changes as input is entered in the data input section, translating static GUI elements into the necessary command line arguments/flags enabling users to view how ChIP-AP’s input is modified based on the provided input.

To instantiate a ChIP-AP run, all input arguments are passed through the command line or the graphical interfaces. What is essential for a run is the location of the input sequencing files (raw fastq) and a settings table for customization of pipeline constituent programs (discussed in the following section) (**Figure 3b**). ChIP-AP then proceeds to then construct a folder hierarchy and places within each folder the corresponding sub-script for that stage of analysis. Each ChIP-AP sub-script script is in essence an instantiated object with defined input and output parameters passing files sequentially from one folder to the next for processing and analysis. Should a user wish to add to or remove an aspect of the pipeline, one simply needs to be mindful of the adjoining objects input/output characteristics. ChIP-AP therefore provides an analysis platform wherein individual analysis steps can be modularly swapped with equivalent steps, provided they have identical input and output characteristics, without requiring additional changes to the flow of analysis or code. This compartmentalization of analytical steps enables ChIP-AP to be exponentially customizable to differing scenarios if the user is proficient enough to code the equivalent analysis step required. The ChIP-AP documentation on GitHub accurately outlines all the analysis steps and documents the input and output behaviours of each sub-script, this is in addition to a comprehensively commented master script outlining the same information in code.

#### Constituent Program Customization and Analysis Reproducibility Through the Settings Table

The lack of result reproducibility in science is a major and on-going issue^35^. The field has continued to change and adapt to this problem with journals enforcing stricter reporting of materials and methods in an attempt to curtail such issues. Unfortunately, bioinformatics methods reporting is an area of scientific research where significant work is still required. Reporting of ChIP-Seq analyses in publications consistently lacks necessary details with many authors omitting key program modification parameters or even neglecting to mention key analysis steps entirely. We have therefore attempted to address this issue by ensuring ChIP-AP analyses are reproducible through an accurate and consistent means of reporting.

A key design aspect of ChIP-AP was to require the provision of a Settings Table (ST). If no table is provided, ChIP-AP will use a pre-defined default-ST (DST; **Table 2**). The ST lists all the programs used in the ChIP-AP run and all the necessary optional program arguments entered for that particular run. It is therefore a listing of all non-hard-coded program modification parameters/flags used for a particular analysis. For ChIP-AP to reproduce any analysis, it simply needs the raw sequencing fastq files and the ST used. *We consider the dissemination of the information contained in the ST as both vital and essential, along with results obtained*. The ST can be included as a supplemental table in a manuscript or can be included as a processed data file when submitting data to an upload repository like GEO. In either case, the information of this file *must* be presented when publishing data to ensure analysis reproducibility in a format that is both consistent and convenient. Of note also, whether the user provided a ST as input or the default-ST was used, a copy of the table will be found in the output folder to ensure all required program modification parameters are provided accompanying the final result.

#### ChIP-AP User Interfaces for Biologists

There is an ever-increasing need to make dry-lab analyses accessible to wet-lab biologists wishing to investigate and interrogate data themselves without having to collaborate with (or wait for) a bioinformatician. This is straightforward if a single program is required for an analysis like GraphPad Prism or SAS. However, longer or more comprehensive analyses and workflows would typically require a degree of coding to work. It is in this domain that ChIP-Seq analysis resides as it requires the utilization of multiple programs, each feeding into each other to perform a coherent analysis. In an attempt to address this demand, platforms such as Galaxy, or licensed software such as the Partek Genomics Suite, have been developed to add graphical user interface (GUI) elements to analyses to make higher-level analyses more accessible to researchers with no coding background. These platforms though, particularly for ChIP-Seq analyses, utilize only a single peak caller and can offer limited customization of program parameters in certain scenarios. As discussed, this can result in incomplete analysis of the bound landscape owing to algorithmic limitations and biases, issues ChIP-AP was designed to address. It was therefore necessary for ChIP-AP to incorporate its own GUI to aide users in completing their required analyses and thus enable researchers with no coding experience to perform independent analyses.

To address the breadth of computer proficiencies seen in the wet-lab scientific community, we implemented two GUI’s, the choice of which to use will depend on a user’s proficiency with ChIP-AP. Through the guided step-by-step tutorials found on our Github, users can install ChIP-AP on any modern operating system, including Windows 10, and run the GUI of their choosing. The GitHub repository lists the system hardware requirements to run ChIP-AP, but many modern laptops and computers commonly purchased for research are capable of analysing data locally, without needing dedicated server hardware.

For users unfamiliar with the command line, we have implemented the Wizard interface (**Figure 3c**), inspired by installation wizards from the Windows 95/98 era of computing. The ChIP-AP wizard will guide users through the analysis configuration by means of a series of panels each asking a single question about the input data. On completion, users will have provided all the necessary information required for a ChIP-AP run and can start their analysis directly from the wizard. This GUI implementation was designed to not overwhelm users with multiple questions simultaneously asking for input, but rather asks for data in a more guided approach.

For users familiar with the input requirements of ChIP-AP, we have implemented the Dashboard interface (**Figure 3d**). The dashboard asks the same questions as the wizard but in a single panel, enabling users to input the required data more quickly (**Figure 3d** – Data Input). Once all the required information is input, as with the wizard, users can run ChIP-AP directly from the interface. In stark difference to the wizard though, the dashboard interface contains a command line translation window at the bottom of the interface (**Figure 3d** – Command Line Translation). As users enter data in the GUI elements, the command line translation window will automatically update to accommodate the additional/changed inputs. This enables researchers to gradually draw connections between translating static GUI elements into command line arguments and flags to modulate and control program behaviour. Such an implementation will aide some researchers more comfortably and confidently transition from GUI to command line usage of ChIP-AP, and hopefully, beyond for their research.

Finally, independent of whether a user opts to use the wizard or dashboard interface, users will be prompted to either use the DST or choose to upload their own ST. As discussed, the functionality and reproducibility provided by the DST/ST is essential for ChIP-AP reproducibility, thus enabling a GUI utilizing researcher to reproduce an analysis performed and customized by a bioinformatician.

### Conclusion

ChIP-Seq is a well-established experimental protocol for profiling DNA-interacting proteins. In the computational space, many software tools have been developed with over 50 peak callers being published to date. Despite the abundance of available peak callers however, benchmarking studies have consistently shown poor overlap between peak-sets from different peak callers. This is because every caller has distinct selectivity and specificity characteristics which are often not additive and seldom completely overlap with other peak callers in many scenarios. Additionally, it has been extensively shown that the performance of a peak caller is subject to the read characteristics and read distributions of the dataset in question. An individual peak caller can outperform other callers in certain datasets but will perform poorly in alternate datasets. Therefore, with the heterogeneity observed in experimental samples arising from profiling different DNA-binding proteins each profiled with differing immunoprecipitation and library preparation protocols, reliance on a single peak caller is unlikely to yield the most reliable, consistent or comprehensive results.

To circumvent the limitations and biases of individual peak callers, we rationalized that integrating the results of multiple peak callers would yield improved peak calling consistency, confidence and more comprehensively assess the binding landscape without requiring additional wet-lab observations. As such, we developed the integrated ChIP-Seq analysis pipeline, ChIP-AP, which takes design decisions from established workflows such as those utilized in consortia projects like ENCODE^18^. ChIP-AP has been coded from the ground-up to be as simple to use as possible for users inexperienced with the command line by providing two GUI’s for use, the wizard or dashboard. ChIP-AP still however remains exponentially customizable for advanced of users by facilitating fine-grained customization of constituent programs through the ST, or, through its provision of customizable modular framework that enables swapping of analysis stages to tailor ChIP-AP for custom workflows. While ChIP-AP has been designed and written specifically for ChIP-Seq analysis, the framework and design principles on which its coded, facilitate its adaptation and use for other existing (ATAC-Seq^36,37^, RIP-Seq^38^, Cut&Run^34^) and future emerging technologies. Should a new, peak caller or analytical tool be developed, minimal changes are required to add an additional step in the pipeline to accommodate the inclusion of said tool. This allows ChIP-AP to be easily modified to work with emerging techniques and any tools that will be specifically developed for such a technique. ChIP-AP can therefore be expanded or enhanced to suit future applications and uses with necessary program arguments being passed through the settings table for each ChIP-AP run. To the best of our research, we have yet to find an integrated software solution currently available that utilizes multiple peak callers other than ChIP-AP.

ChIP-AP as presented here though, allows users to sub-set the binding landscape in a manner that is best suited to address their biological research question, while allowing users to switch between differing sub-sets depending on the question at hand. By utilizing the consensus peak set, binding motif accuracy can be significantly increased by restricting the motif search space to only the most confident peaks. This also has improved outcomes when attempting down-stream GO analysis wherein more targeted and biologically significant terms can be reported. In contrast though, if the profiled dataset has unfavourable characteristics such as poor enrichment or shows high signal:noise, the union peak set can potentially yield improved results and allow users to marginally sacrifice specificity for a potentially significant increase in sensitivity across the binding landscape. In between these two extremes of data sub-sets is a gradient of sensitivity thresholds that can be selected depending on the biological question and the presence of additional, supportive data from independent techniques and methodologies. By reporting such an integrated analysis, ChIP-AP enables the end user to focus on the biological question at hand by providing a comprehensive protein binding profile without needing data re-analysis. ChIP-AP can therefore provide both substantial improvements to peak capturing and analysis reliability from a single integrated and comprehensive analysis.

## Materials and Methods

### ChIP-AP Constituent Programs

ChIP-AP is an integrated pipeline that brings multiple command line programs together into a single, seamless and easy to use pipeline. At time of publication, these include FastQC^39^, Clumpify and BBDuk from the BBMap Suite^40^, Trimmomatic^41^, BWA^42^, Samtools^43^, deepTools^44^, MACS2^45^, GEM^16^, SICER2^46^, HOMER^15^ and Genrich^13^. If using ChIP-AP, please cite all constituent tools as well. It is best to refer to the GitHub repository for the latest citation list which would include any additional tools incorporated into ChIP-AP since publication.

### SNU-398 Culturing

SNU-398 cell line was obtained from the American Type Culture Collection (ATCC). The cells were maintained in RPMI medium supplemented with 10% foetal bovine serum (FBS) at 37°C in a humidified atmosphere of 5% CO2 as recommended by ATCC.

### SALL4 ChIP-Seq Preparation and Sequencing

20 million SNU-398 cells were cross-linked with 1% formaldehyde for 10 minutes at room temperature. The reaction was terminated by adding 2M glycine to a final concentration of 125mM. Cells were then washed with 1×PBS and resuspended in 1mL of cell lysis buffer (20mM Tris pH8.0, 85mM KCl, 0.5% nonidet P-40, protease inhibitor). After 10 minutes of incubation on ice, cells were spun down and cell pellet was resuspended in another 1mL of cell lysis buffer. After another 5 minutes of incubation on ice, cells were spun down and cell pellet was resuspended in 1mL of nuclear lysis buffer (10mM Tris-HCl pH7.5, 1% nonidet P-40, 0.5% sodium deoxycholate, 0.1% SDS, protease inhibitor). After 10 minutes of incubation on ice, chromatin was sheared to 500bp. Antibody-protein A/G Dynabead conjugate was prepared by adding 0.75μg of SALL4 rabbit monoclonal antibody (Cell Signaling Technology #8459) to pre-washed 50μL of protein A/G Dynabeads (Life Techonlogies) with one hour incubation at 4°C with rotation. Sheared chromatin was then added to antibody-protein A/G conjugate and incubated overnight at 4°C with rotation. After overnight incubation, the beads were washed sequentially with the following buffers: twice with RIPA/500mM NaCl buffer (0.1% deoxycholate, 0.1% SDS, 1% Triton X-100, 500mM NaCl, 1mM EDTA, 20mM Tris-HCl pH8.1), twice with LiCl buffer (0.25M LiCl, 1% nonidet P-40, 1% sodium deoxycholate, 1mM EDTA, 10mM Tris-HCl pH8.1), twice with TE buffer (10mM Tris-HCl pH8.0, 1mM EDTA pH8.0). Protein complexes were reverse cross-linked with 50μL of ChIP Elution Buffer (10mM Tris-HCl pH8.0, 5mM EDTA, 300mM NaCl, 0.1% SDS) and 8μL of Reverse Crosslink Mix (250mM Tris-HCl pH6.5, 1.25M NaCl, 62.5mM EDTA, 5mg/mL proteinase K, 62.5μg/mL RNase A) at 65°C for 5 hours. Reverse cross-linked DNA was cleaned up using SPRI beads (Beckman Coulter) and eluted in 10mM Tris-HCl pH 8.0. To generate libraries for deep sequencing, the eluted DNA was end-repaired using End-It DNA End-Repair Kit (Epicenter #ER0720) and A-tailing was then carried using Klenow (3’-5’ exo-) enzyme (New England Biolabs). Illumina sequencing adaptors were ligated to the DNA fragments and adaptor-ligated DNA fragments were enriched with 14 cycles of PCR. DNA libraries were gel purified and analyzed on Bioanalyzer (Agilent) for their size distribution. Libraries were sequenced on Illumina HiSeq 2500 sequencer with single-end 35bp settings. The sequencing and processed files have been uploaded to GEO with Accession number xxxx (reviewer access token xxxx).

### SALL4 ChIP-Seq Analysis and Comparisons

The generated SALL4 ChIP-Seq was processed with ChIP-AP (v4.1) using a hg38 genome. The settings table used for analysis is found below.

| Program | Argument |
| --- | --- |
| fastqc1 | -q |
| clumpify | dedupe spany addcount |
| bbduk | ktrim=1 hdist=2 |
| trimmomatic | LEADING:20 SLIDINGWINDOW:4:20 TRAILING:20<br>MINLEN:20 |
| fastqc2 | -q |
| bwa_mem |  |
| samtools_view |  |
| plotfingerprint | -bs 50 --centerReads --ignoreDuplicates |
| fastqcs3 | -q |
| macs2_callpeak |  |
| gem | -Xmx30G --k_min 8 --k_max 12 |
| sicer2 |  |
| homer_findPeaks |  |
| genrich | --adjustp -v |
| homer_mergePeaks |  |
| homer_annotatePeaks |  |
| fold_change_calculator | --normfactor uniquely_mapped |

For all analyses, the union peak set was utilized. The fingerprint plot (Figure 2a) was generated as part of the ChIP-AP run with the flags outlined in the settings table. The upset plot (Figure 2b) was generated by taking the “venn.txt” data from the ChIP-AP run output (folder 21_peaks_merging) and plotting it in R^47^ (v4.0.3) with the UpSetR^48^ (v1.4.0) package.

For comparisons with the Cut&Run data, the Cut&Run data was processed as outlined previously^33^ and is available from GEO, Accession GSE136332. To overlap the Cut&Run replicates, HOMER’s mergePeaks was used with flags “-d 1500.” Next, the Cut&Run peaks identified in at least 2 replicates were combined into a single list and compared to the SALL4 ChIP-Seq union peak set using HOMER’s mergePeaks with flag “-d 2000”, this provided the list of overlapping regions, the number of which was plotted in R^47^ (v4.0.3) with the VennDiagram^49^ (v1.6.20) package.

For the directed motif search within the SALL4 ChIP-Seq union peak set, HOMER’s^15^ findMotifsGenome function was used with flags “-find sall4_weighted_motif.motif.” For the HOMER *de novo* motif search, HOMER’s findMotifsGenome function was used with flags “hg38 - size given -mask.” For the MEME-ChIP (and sub-program^22–24^) motif search, first the union peak list was processed with HOMER’s findMotifsGenome function with flag “-dumpFasta” to extract the central 200bp sequences of each peak. HOMER also generated an equivalent set of background sequences with comparable GC content to be used. Next, MEME-ChIP was run with flags “-neg background.fa -meme-nmotifs 25 union_peaks.fa.” Motif logo files were generated using R^47^ (v4.0.3) and the seqLogo^50^ (v1.52.0) package.

The gene ontology analysis of the SALL4 ChIP-Seq dataset performed was part of the ChIP-AP run using the flag “-goann” which utilizes HOMER to perform the analysis following peak annotation. To compare with the processed SALL4 knock-down results published^33^, we started from supplemental tables 4 and 5 from the publication. Next, we overlayed the reported gene names from the SALL4 ChIP-Seq union peak set to those gene lists to determine overlapping gene names.

### Encode Datasets Utilized and Processing

A number of ENCODE datasets were downloaded and utilized for our analysis. The table below lists all the downloaded experiment ID’s used. Data was downloaded from ENCODE March 2021.

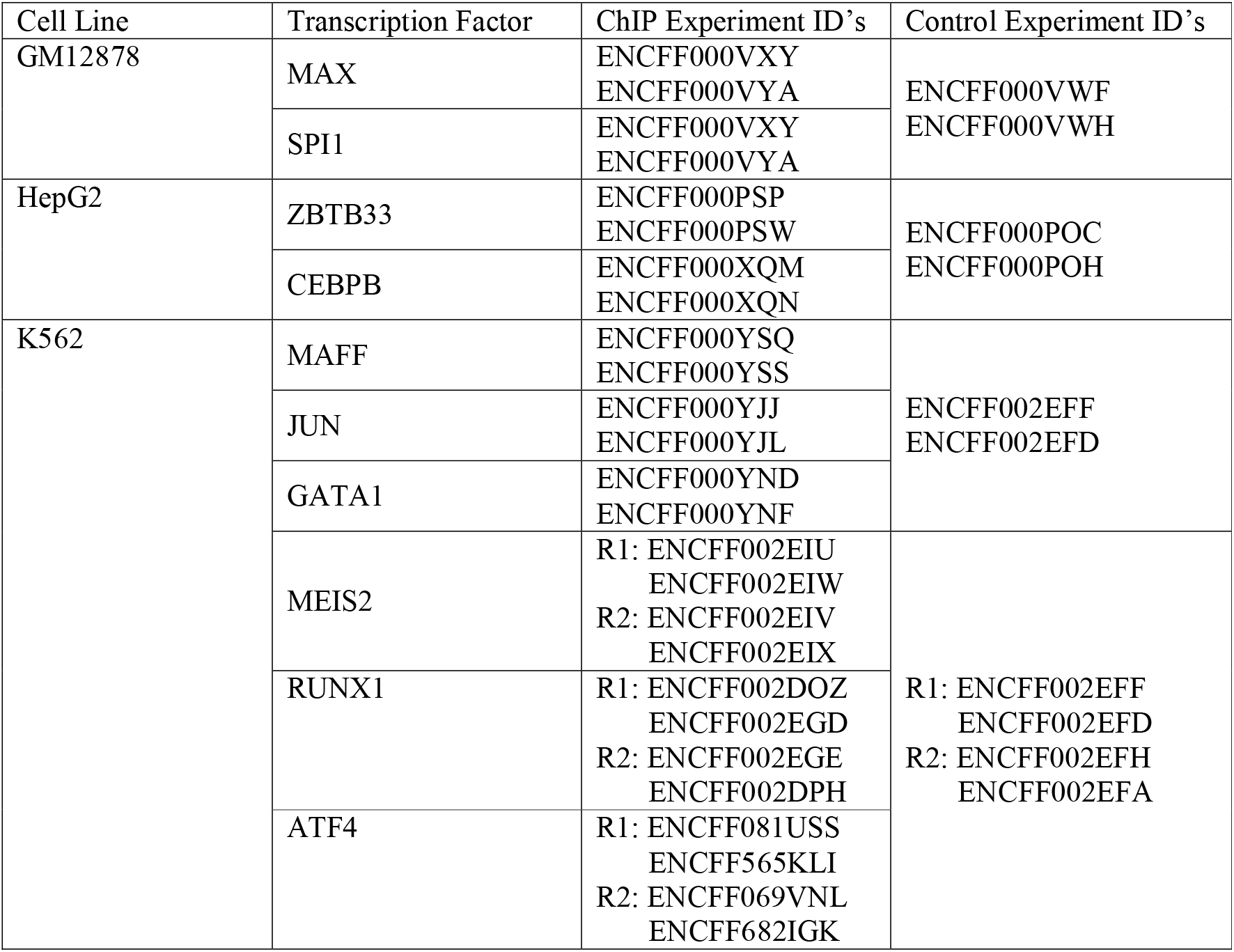

| Cell Line | Transcription Factor | ChIP Experiment ID's | Control Experiment ID's |
| --- | --- | --- | --- |
| GM12878 | MAX | ENCFF000VXY<br>ENCFF000VYA | ENCFF000VWF<br>ENCFF000VWH |
|  | SPI1 | ENCFF000VXY<br>ENCFF000VYA |  |
| HepG2 | ZBTB33 | ENCFF000PSP<br>ENCFF000PSW | ENCFF000POC<br>ENCFF000POH |
|  | CEBPB | ENCFF000XQM<br>ENCFF000XQN |  |
| K562 | MAFF | ENCFF000YSQ<br>ENCFF000YSS | ENCFF002EFF<br>ENCFF002EFD |
|  | JUN | ENCFF000YJJ<br>ENCFF000YJL |  |
|  | GATA1 | ENCFF000YND<br>ENCFF000YNF |  |
|  | MEIS2 | R1: ENCFF002EIU<br>ENCFF002EIW<br>R2: ENCFF002EIV<br>ENCFF002EIX | R1: ENCFF002EFF<br>ENCFF002EFD<br>R2: ENCFF002EFH<br>ENCFF002EFA |
|  | RUNX1 | R1: ENCFF002DOZ<br>ENCFF002EGD<br>R2: ENCFF002EGE<br>ENCFF002DPH |  |
|  | ATF4 | R1: ENCFF081USS<br>ENCFF565KLI<br>R2: ENCFF069VNL<br>ENCFF682IGK |  |

All ENCODE datasets were processed with ChIP-AP (v4.1) with the following settings table

| Program | Argument |
| --- | --- |
| fastqc1 | -q |
| clumpify | dedupe spany addcount qout=33 fixjunk |
| bbduk | ktrim=1 hdist=2 |
| trimmomatic | LEADING:20 SLIDINGWINDOW:4:20 TRAILING:20 MINLEN:20 |
| fastqc2 | -q |
| bwa_mem |  |
| samtools_view |  |
| plotfingerprint | -bs 50 --centerReads --ignoreDuplicates |
| fastqcs3 | -q |
| macs2_callpeak |  |
| gem | -Xmx30G --k_min 8 --k_max 12 |
| sicer2 |  |
| homer_findPeaks |  |
| genrich | --adjustp -v |

For each transcription factor, the corresponding JASPAR binding motif for the cell line in question was downloaded from MethMotif^19^ and manually converted to HOMER motif format. For the directed motif searches, HOMER’s findMotifsGenome was utilized with flags “hg38 -find binding_motif.motif.” For HOMER *de novo* motif discovery, the findMotifsGenome program was used with flags “hg38 -size given -mask -dumpFasta.” This ran the motif discovery while also giving the necessary fasta sequence files (target.fa and background.fa) required to run the MEME-Suite. The MEME *de novo* motif discovery was run with flags “-neg background.fa -meme-nmotifs 25 target.fa.” Motif logo files were generated using R^47^ (v4.0.3) and the seqLogo^50^ (v1.52.0) package. The gene ontology results were generated as part of HOMER’s annotatePeaks function for the required peak sets. HOMER annotatePeaks was utilized with a known motif provided with -m flag to include the distances from all starting coordinate motif instances in each peak to their respective peak starting coordinate. A custom script was utilized to extract the distances from every peak’s weighted peak center coordinate to the midpoint coordinate of the motif instance closest to the weighted peak center. The density plots representing this data were generated using R^47^ and the ggplot2^51^. Peak-Motif percentages were plotted using Graphpad Prism v9.1.0.

---

Supplemental Figure 1

a) The motif position-bias for ATF4, GATA1, MAFF, MAX, MEIS2 and RUNX1 for the consensus peak set and the individual peak callers. The position-bias is a measure of how far the identified motif sits away from the weighted peak center. b) The CEBPB *de novo* motif discovery results as reported by the MEME-Suite for the consensus peak set. Above each p-value is which sub-program of MEME called the said motif. c) The CEBPB *de novo* motif discovery results as reported by the MEME-Suite for the MACS2 peak set. Above each p-value is which sub-program of MEME called said motif. d) The MAFF *de novo* motif discovery results as reported by the MEME-Suite for the consensus peak set. Above each p-value is which sub-program of MEME called said motif. The 6^th^ result shows a characteristic heterodimer binding profile for MAFF. e) The MAFF *de novo* motif discovery results as reported by the MEME-Suite for the MACS2 peak set. Above each p-value is which sub-program of MEME called said motif.

Supplemental Figure 2

a) Venn diagrams highlighting the degree of overlap between each individual callers peak-set and the Cut&Run peak set, and the relatively few peaks called by each individual peak caller. b) The MEME-ChIP results highlighting showing the correct AT-rich binding motif for SALL4 is the 3^rd^ called motif hit. The 2^nd^ motif hit also is an AT-rich motif with near identical sequence.

Table 1 – Top 20 RUNX1 GO Terms

The top 20 RUNX1 GO terms returned for the consensus peak set (left) and the MACS2 peak set (right). The consensus peak set GO returned GO terms are more directly relatable to defined RUNX1 functions as compared to the MACS2 results.

Table 2 – Default Settings Table for ChIP-AP

The default program settings table used by ChIP-AP if no user provided settings table is provided. The left column lists the constituent programs of ChIP-AP with their optional modification parameters/flags found in the right column.

Supplemental Table 1

Table listing and overview of all the profiled TF’s, their TF family and cell line of origin. The table also lists the total number of peaks found in the MACS2 and consensus peak sets, along with the peak-motif percentages for each set.

Supplemental Table 2

A listing of the z-tests performed testing the position-bias distributions of the consensus peak set compared to each individual peak caller. Significant differences are highlighted in green. Cells highlighted yellow indicate values approaching significance.

Supplemental Table 3

All the consensus peak GO results for all 10 TF’s (1 sheet per TF).

Supplemental Table 4

All the MACS2 peak GO results for all 10 TF’s (1 sheet per TF).

Supplemental Table 5

The union peak-list for the SALL4 ChIP-seq

Supplemental Table 6

The peak-motif percentages for each individual peak callers results in the SALL4 ChIP-Seq dataset.

Supplemental Table 7

The GO results for the SALL4 union peak-set.

Supplemental Table 8

The differentially expressed genes from the SALL4 knock-down RNA-Seq experiment found to contain at least 1 peak in the SALL4 union peak-set.

## Acknowledgements

The authors would like to thank Yanjing V. Liu, Giorgia Maroni, Elena Levantini, Chenxi Qiu and Bon Q. Trinh for their feedback on the ChIP-AP installation guides and installations. We want to thank them for letting us use their laptops and computers as test-beds to ensure our software is as seamless as possible for others to use. We also want to thank Robert Welner for his feedback on the manuscript. We want to especially thank Touati Benoukraf for his insightful discussions and comments regarding ChIP-AP, its development and its presentation.

## Contributions

J.S. and M.A.B designed the package. J.S. was the lead programmer. J.S. and M.A.B tested, optimized and debugged ChIP-AP. D.E.T and KJ.Y. performed the SALL4 ChIP-Seq. J.S., D.G.T and M.A.B interpreted results and wrote the manuscript. M.A.B conceived and directed the project.

