## Supplementary material for "ChIP-AP – An Integrated ChIP-Seq Analysis Pipeline": Table 1

| Consensus Peak Set Gene Ontology (Biological Processes (Results) | | | | MACS2 Peak Set Gene Ontology (Biological Processes (Results) | | | |
| --- | --- | --- | --- | --- | --- | --- | --- |
| TermID | **Term** | **Enrichment** | **Target Genes  in Term** | **TermID** | **Term** | **Enrichment** | **Target Genes  in Term** |
| GO:0065009 | regulation of molecular function | 8.44*10^-07^ | 82 | **GO:0048518** | positive regulation of biological process | 3.59*10^-13^ | 935 |
| GO:0010506 | regulation of autophagy | 3.38*10^-05^ | 17 | **GO:0048522** | positive regulation of cellular process | 6.37*10^-13^ | 832 |
| GO:0032870 | cellular response to hormone stimulus | 5.63*10^-05^ | 24 | **GO:0071840** | cellular component organization or biogenesis | 2.25*10^-12^ | 880 |
| GO:0051716 | cellular response to stimulus | 8.93*10^-05^ | 140 | **GO:0031325** | positive regulation of cellular metabolic process | 2.49*10^-12^ | 540 |
| GO:2000973 | regulation of pro-B cell differentiation | 0.000144 | 3 | **GO:0065009** | regulation of molecular function | 4.77*10^-12^ | 493 |
| GO:1900221 | regulation of amyloid-beta clearance | 0.000178 | 4 | **GO:0031323** | regulation of cellular metabolic process | 5.05*10^-12^ | 938 |
| GO:0070887 | cellular response to chemical stimulus | 0.000185 | 71 | **GO:0016043** | cellular component organization | 5.93*10^-12^ | 852 |
| GO:1905456 | regulation of lymphoid  progenitor cell differentiation | 0.000227 | 3 | **GO:0009893** | positive regulation of metabolic process | 7.84*10^-12^ | 581 |
| GO:0090218 | positive regulation of lipid kinase activity | 0.00028 | 5 | **GO:0043412** | macromolecule modification | 1.05*10^-11^ | 529 |
| GO:1903432 | regulation of TORC1 signaling | 0.000319 | 5 | **GO:0010604** | positive regulation of macromolecule  metabolic process | 1.06*10^-11^ | 542 |
| GO:0007163 | establishment or maintenance of cell polarity | 0.000329 | 11 | **GO:0036211** | protein modification process | 1.42*10^-11^ | 499 |
| GO:1901700 | response to oxygen-containing compound | 0.000336 | 43 | **GO:0006464** | cellular protein modification process | 1.42*10^-11^ | 499 |
| GO:0050790 | regulation of catalytic activity | 0.000376 | 59 | **GO:0051173** | positive regulation of nitrogen compound metabolic process | 2.19*10^-11^ | 514 |
| GO:0071495 | cellular response to endogenous stimulus | 0.000391 | 35 | **GO:0050790** | regulation of catalytic activity | 3.36*10^-11^ | 394 |
| GO:1901699 | cellular response to nitrogen compound | 0.000405 | 22 | **GO:0044260** | cellular macromolecule metabolic process | 3.55*10^-11^ | 765 |
| GO:0019216 | regulation of lipid metabolic process | 0.00044 | 17 | **GO:0048523** | negative regulation of cellular process | 3.88*10^-11^ | 739 |
| GO:1900222 | negative regulation of amyloid-beta clearance | 0.000475 | 3 | **GO:0006996** | organelle organization | 5.43*10^-10^ | 534 |
| GO:0016241 | regulation of macroautophagy | 0.000519 | 10 | **GO:0051128** | regulation of cellular component organization | 8.46*10^-10^ | 406 |
| GO:0044093 | positive regulation of molecular function | 0.000575 | 47 | **GO:0031329** | regulation of cellular catabolic process | 8.97*10^-10^ | 164 |
