## Supplementary material for "ChIP-AP – An Integrated ChIP-Seq Analysis Pipeline": Table 2

| program | argument |
| --- | --- |
| fastqc1 | -q |
| clumpify | dedupe spany addcount qout=33 fixjunk |
| bbduk | ktrim=l hdist=2 |
| trimmomatic | LEADING:20 SLIDINGWINDOW:4:20 TRAILING:20 MINLEN:20 |
| fastqc2 | -q |
| bwa_mem |  |
| samtools_view | -q 20 |
| plotfingerprint |  |
| fastqc3 | -q |
| macs2_callpeak |  |
| gem | -Xmx10G --k_min 8 --k_max 12 |
| sicer2 |  |
| homer_findPeaks |  |
| genrich | --adjustp -v |
| homer_mergePeaks |  |
| homer_annotatePeaks |  |
| fold_change_calculator | --normfactor uniquely_mapped |
