## Supplementary figures and images for "ChIP-AP – An Integrated ChIP-Seq Analysis Pipeline"

### Supplemental Figures

#### Supplemental Figure 1

**a**

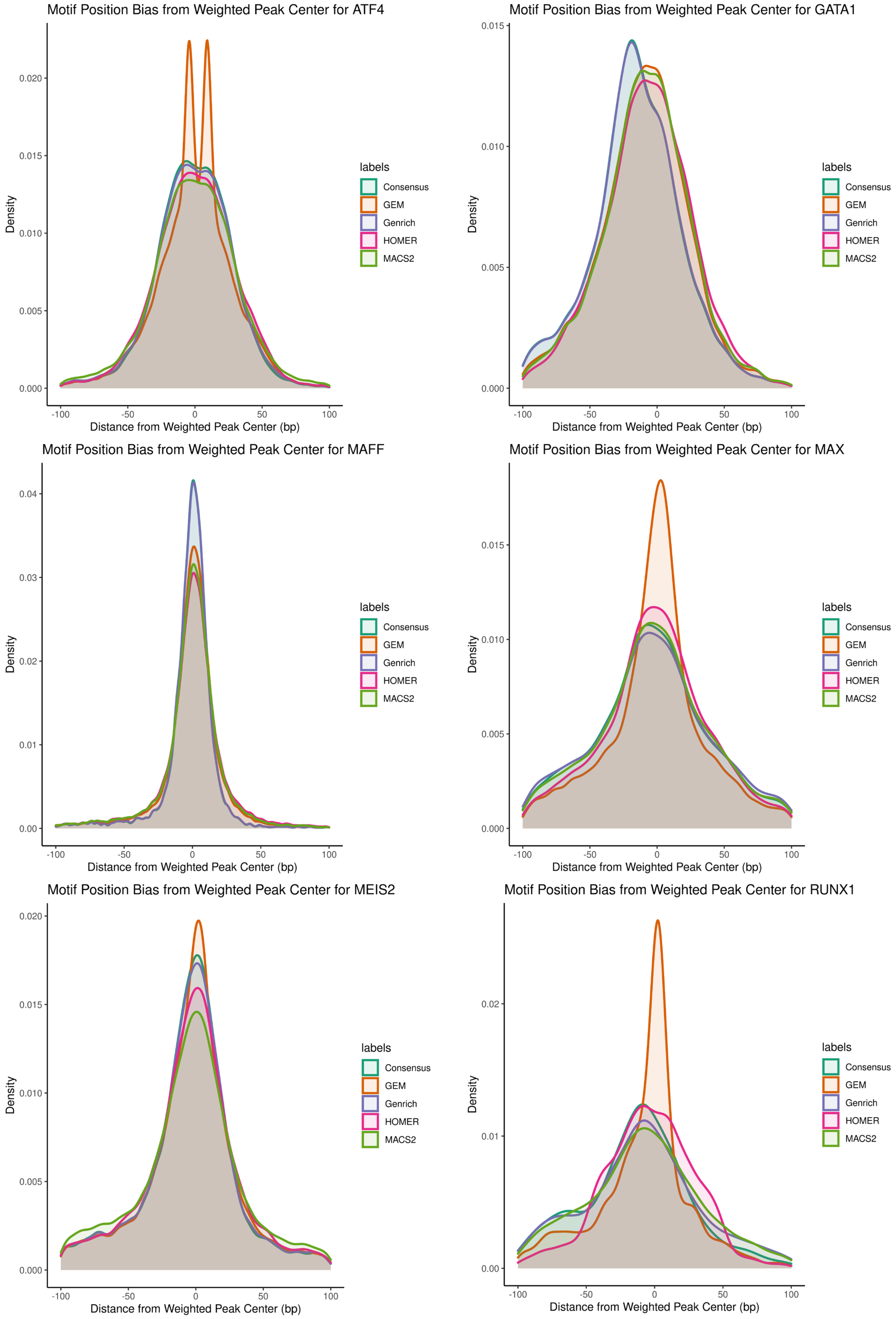

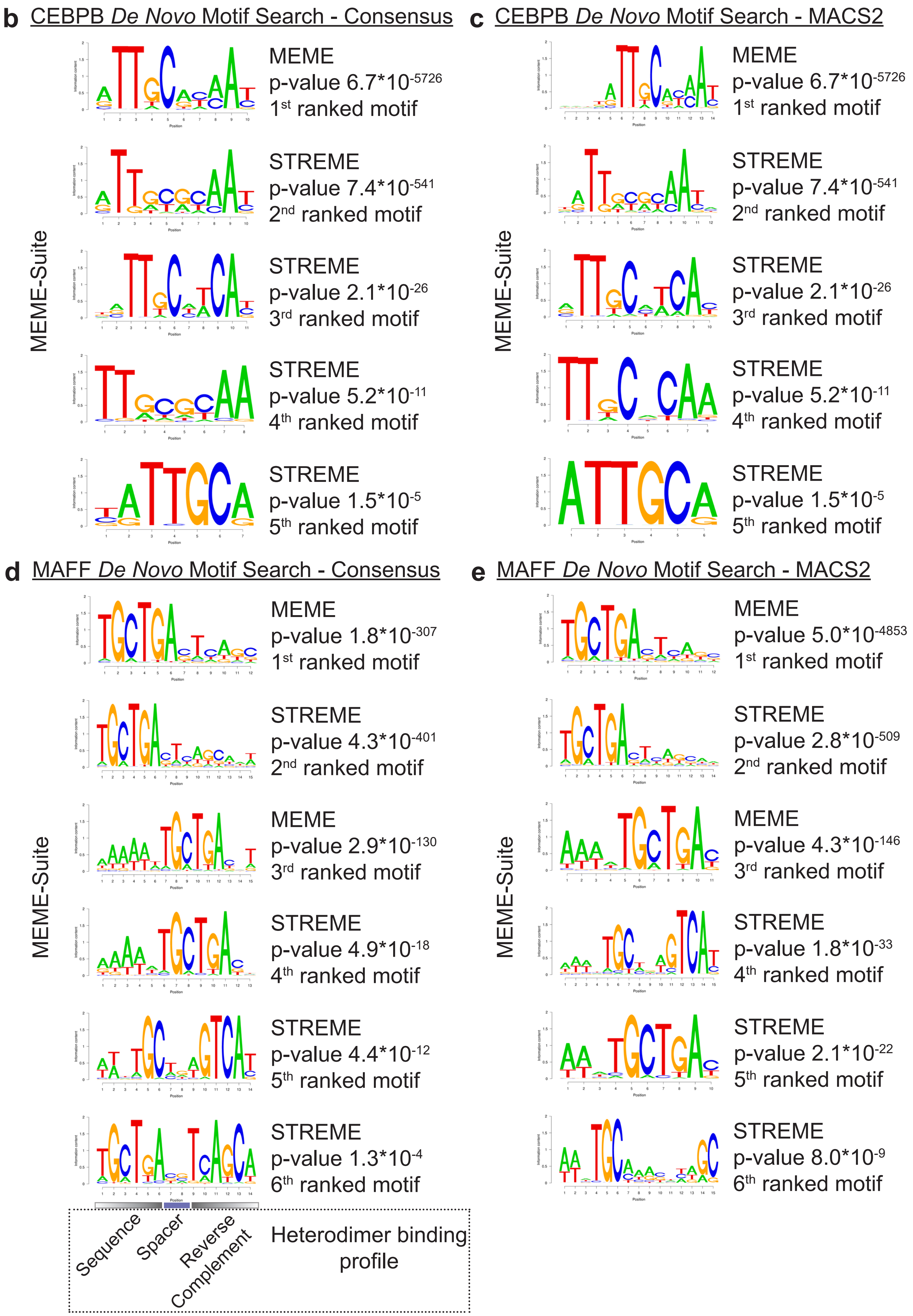

#### Supplemental Figure 2

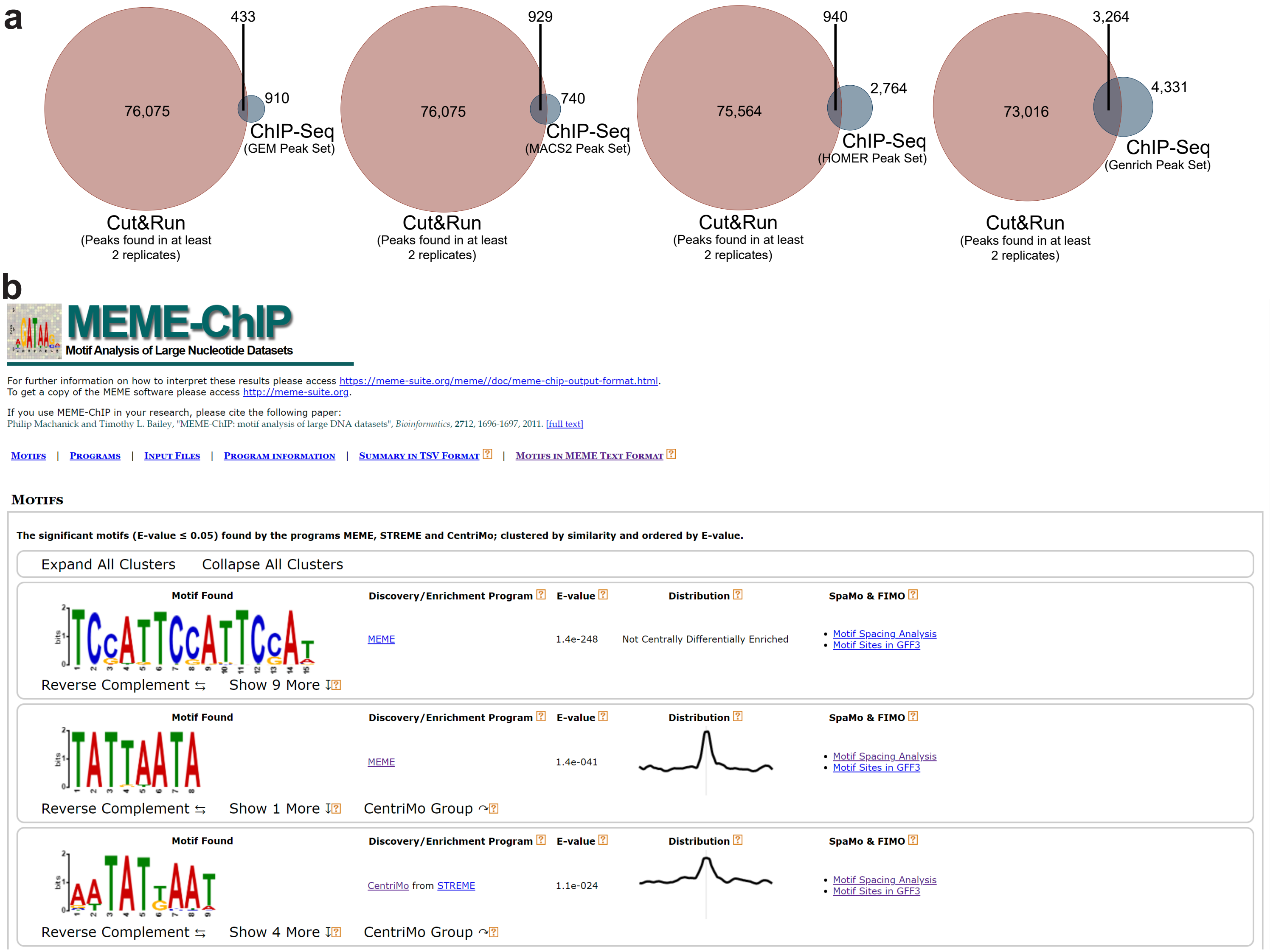
